# Seahorse Metabolic Analysis for Human and Mouse Cardiac Organotypic Slices

**DOI:** 10.1101/2025.06.11.659150

**Authors:** Katy A Trampel, Binjie Li, Altynai Melisova, Micah Madrid, Sharon A George, Igor R Efimov

## Abstract

The mammalian heart relies on high rates of mitochondrial oxidative phosphorylation to meet its energy demand, with fatty acids serving as the primary fuel source in healthy adult hearts. While metabolic flexibility, the ability to switch between metabolic fuel substrates, is known to change during development and cardiac diseases, standardized methods for assessing substrate usage in intact, living cardiac tissue remain limited. Here, we present a protocol that adapts the Seahorse Mito Fuel Flex Test for use in living organotypic cardiac slices. This method enables the quantification of fuel dependency and capacity for fatty acids (FA), glucose (GLC), and glutamine (GLN) by sequentially inhibiting their respective mitochondrial oxidative phosphorylation pathways with the inhibitors etomoxir, UK5099, and BPTES. First, we validated the protocol by comparing results from organotypic cardiac slices to the standard published protocol using isolated adult mouse primary cardiomyocytes. Next, we demonstrated the sensitivity of this assay by modulating metabolism with AICAR, an AMPK activator, at varying concentrations, to demonstrate improved metabolism and then metabolic suppression at higher toxic doses. Finally, we applied this protocol to organotypic cardiac slices from different chambers of human donor hearts. This protocol provides a high-throughput, physiologically relevant platform for investigating cardiac metabolism, applicable across species and adaptable to other tissue types. It enables the study of metabolic remodeling in development and disease while overcoming the limitations of traditional cell-based assays by preserving native tissue architecture, physiology, and multicellular heterogeneity.

## INTRODUCTION

The heart is a highly energetic organ that continuously hydrolyzes adenosine triphosphate (ATP) to sustain its contractile and other functions. Approximately 60 – 70% of ATP hydrolysis in the heart fuels contraction, while 30 – 40% powers ion pumps, such as the sarcoplasmic reticulum Ca²⁺-ATPase (SERCA)^1–3^. Under non-ischemic conditions, a healthy human heart produces greater than 95% of its ATP through oxidative phosphorylation in the mitochondria, with the remainder produced via glycolysis and GTP synthesis in the tricarboxylic acid (TCA) cycle. To meet its high energy demands, the heart metabolizes various energy substrates, including triacylglycerols, long-chain fatty acids (FA), glucose (GLC), glycogen, lactate, pyruvate, ketone bodies, and amino acids (especially branch-chained amino acids). These substrates are catabolized into TCA cycle intermediates and electron transport carriers (NADH and FADH_2_), which fuel oxidative phosphorylation^4^. Fatty acids serve as a primary energy source, providing 60 – 90% of cardiac ATP through β-oxidation, while carbohydrates contribute 10 – 40% through glycolysis and GLC oxidation^1,5^.

In contrast to healthy adult hearts, fetal and pathological hearts exhibit altered metabolic substrate preferences. Developing hearts primarily rely on carbohydrates to fuel glycolysis and lactate oxidation, enabling ATP production under low oxygen conditions (physiologic hypoxia)^6^. Similarly, in pathological states such as pressure overload, hypertrophy, and ischemia, metabolism shifts toward glycolysis and GLC oxidation to enhance oxygen efficiency for ATP synthesis during hypoxic conditions^1,7^.

The Seahorse Mito Fuel Flex Test is a metabolic assay designed to evaluate fuel utilization by systemically inhibiting key metabolic pathways for glucose (GLC, pyruvate), glutamine (GLN, glutamate), and long-chain fatty acids (FAs). This assay quantifies parameters such as fuel dependency and fuel capacity. Fuel dependency is defined as the reliance of cells on a specific fuel source to maintain baseline respiration. Fuel capacity is defined as the ability of cells to utilize a given fuel source when all other fuel pathways are blocked. The interplay between dependency and capacity defines metabolic flexibility, which reflects the cell’s ability to dynamically adjust mitochondrial energy production by shifting between different fuel sources. Oxygen consumption rate (OCR) is measured in response to specific combinations of metabolic inhibitors across six parallel tests performed in different wells of the Seahorse microculture plate. FA dependency is assessed by injecting etomoxir to inhibit FA oxidation, followed by UK5099 and BPTES to inhibit GLC and GLN oxidation, respectively. FA capacity is measured by reversing this injection order. GLC dependency involves initial injection of UK5099, followed by BPTES and etomoxir, while GLC capacity is measured by injecting BPTES and etomoxir first, then UK5099. GLN dependency is tested by injecting BPTES first, followed by etomoxir and UK5099, and GLN capacity is assessed by reversing this order. At the start of the assay, baseline respiration is recorded before inhibitor injections. Etomoxir blocks FA oxidation by inhibiting carnitine palmitoyl-transferase 1A (CPT1A)^4^, UK5099 inhibits mitochondrial pyruvate carrier to block GLC oxidation^8^, and BPTES is an allosteric glutaminase inhibitor that prevents GLN conversion to TCA cycle intermediates^9^. Together, these tests provide a comprehensive profile of mitochondrial fuel preference and metabolic adaptability.

### Development of Protocol

In this study, Seahorse Real-Time Cell Metabolic Analysis, specifically the Seahorse Mito Fuel Flex Test, was adapted for cardiac organotypic slices. Inhibitor-dose response testing was performed to determine the half-maximal inhibitory concentration (IC_50_) of etomoxir, UK5099, and BPTES for organotypic slices, as there are differences in permeability, cell count, and cell type diversity in tissue preparations compared to isolated cell models.

Since FAs are the primary energy source of cardiomyocytes, we optimized the assay by supplementing the media with exogenous FA (Bovine Serum Albumin (BSA)-palmitate) to determine the optimal concentration that best recapitulates cardiomyocyte FA utilization, as the standard Seahorse Mito Fuel Flex Test Media (Agilent Seahorse XF DMEM Medium) lacks FAs. The Seahorse Mito Fuel Flex Test was then performed to assess mitochondrial fuel dependency and capacity for FAs, GLC, and GLN in healthy mouse cardiac organotypic slices. To validate the assay, we compared our results in organotypic slices to the standard Seahorse protocol for isolated primary adult mouse cardiomyocytes. We tested whether Acadesine (AICAR), an AMPK activator, enhances metabolic response in mouse LV cardiac organotypic slices^10^. Finally, we applied this technique to compare metabolism in the four chambers of donor human hearts, extending its relevance to human cardiac metabolism.

### Applications of Methods

This adaptation of the standardized method for Seahorse Metabolic Analysis, specifically the Seahorse Mito Fuel Flex Test, extends its application to various species and organ tissues, broadening its utility beyond cardiac research to neurological diseases, cancer, and aging. This approach serves as a powerful tool for investigating metabolic disruptions in pathologies and understanding developmental metabolic changes. By modifying the media composition to more accurately reflect the physiological metabolite environment with FAs, this method enables the study of cardiac pathophysiology under more biologically relevant conditions. Furthermore, the ability to scale inhibitor concentrations from cell-based Seahorse assays to the levels required for tissue studies allows for a more precise assessment of metabolic regulation across different biological models and, potentially, in the clinic.

### Comparison with Other Methods

Cardiac metabolism has previously been assessed by radionucleotide labeling of energy substrates, such as at ^3^H and ^14^C, to determine substrate utilization *in vivo* via photon emission tomography (PET) and single-photon emission computed tomography (SPECT)^11^. Similarly, ^31^P, ^1^H, and ^13^C nuclear magnetic resonance (NMR) imaging has been employed both *in vivo* and *ex vivo* to measure metabolic flux^12^. While these techniques are highly sensitive and effective in quantifying metabolism under baseline conditions, they are limited by the number of pathways that can be labeled and the technical expertise and equipment required for these techniques.

Metabolic flux was initially assessed by measuring pressure changes in small samples of blood to quantify oxygen consumption^13^. Since then, Seahorse Real-Time Cell Metabolic Analysis has been developed using fluorescence-based technology to measure oxygen levels and pH, corresponding to mitochondrial oxidation and glycolysis, respectively, in live cells *in vitro*. This method offers a high signal-to-noise ratio and is sensitive enough to measure flux in small cell populations and mitochondrial samples. However, 2D cell culture does not fully recapitulate the 3-dimensional architecture, native extracellular matrix, and numerous cell types in the myocardium or other organs and tissues. In the context of cardiac metabolism, isolating adult primary cardiomyocytes is suboptimal, as enzymatic digestion alters their morphology, transcriptome, metabolic processes, and disrupts cell-cell interactions, limiting their physiological relevance for metabolic studies^14–17^.

Despite success in cell-based assays, adapting Seahorse Real-Time Cell Metabolic Analysis to measure metabolic flux in intact tissue remains challenging. Protocols for organoids and cancer spheroids have been refined to enable 3D metabolic studies^18,19^. In neurological research, tissue sectioning has become routine for studying metabolic flux using Seahorse Metabolic Analysis, which creates uniform tissue thickness, and biopsy punches can be used to fit the tissue slice into the Seahorse microculture well plate^20–23^. However, some protocols embed tissue in agarose before sectioning, which can limit oxygen diffusion and affect tissue viability.

Few studies have adapted cell-based Seahorse protocols for intact, living cardiac tissue^24,25^. Existing studies rely on the Seahorse XFe24-analyzer, which has lower throughput than the Seahorse XFe96-analyzer. Additionally, these studies lack standardization, leading to inconsistencies in tissue thickness, media composition, and fuel availability, all of which are crucial for accurately assessing cardiomyocyte fuel utilization. Furthermore, many studies fail to normalize metabolic data to tissue area, as the biopsy punch method often produces imperfect tissue samples. These limitations underscore the need for a refined and standardized protocol to improve the reproducibility and accuracy of living cardiac tissue metabolic analysis using Seahorse Metabolic Analysis. Notably, no tissue slice Seahorse Metabolic Analysis Protocols have been adapted for the Seahorse Mito Fuel Flex Test, which provides insights into fuel substrate utilization, a mechanism known to change during development and cardiac disease.

### Experimental Design

When adapting Seahorse Metabolic Analysis for tissue preparations, several critical experimental design factors must be considered to optimize tissue-based assays. One critical factor is the space available in each well of the XFe96 culture plate, which is limited to a height of 200 μm and a bottom well diameter of 3.81 mm. To ensure adequate space for 180 μL of media and all injection port inhibitor volumes, tissue slices must be thin enough to fit within this confined space. Therefore, we used a 100 μm, 2 mm diameter tissue section in our protocol. This thickness allowed for adequate oxygenation of the tissue.

Next, inhibitor concentrations were adjusted for the tissue-based assay, as the doses used for cell-based Seahorse assays may be insufficient. A preliminary IC_50_ study should be performed to determine the optimal concentration of inhibitors used. Data from the Seahorse Mito Fuel Test in this study can be used to scale inhibitors appropriately for other Seahorse Metabolic Assays. Additionally, the media composition is also crucial for accurate metabolic measurements. If FA, GLC, and GLN dependency and capacity are being assessed, the media must contain FA, as it is the primary energy source of the heart. Agilent Seahorse XF DMEM Medium lacks FA, so BSA-palmitate supplementation was applied to the media to better replicate physiological conditions.

Lastly, to ensure reproducibility, technical replicates and data normalization should be applied as required. We recommend that each experimental group include a minimum of three technical replicates and biological replicates. Data normalization should be performed using tissue area in each well to minimize intersample variability. To perform this normalization, images of each well should be acquired at the end of each protocol to calculate the tissue area. Larger slices contain more cells and thus exhibit higher metabolic rates. Thus, normalizing metabolic data to tissue area is essential for accurate comparisons across wells.

### Expertise required to implement the protocol

Successfully adapting Seahorse Metabolic Analysis for cardiac tissue requires expertise in aortic cannulation, cardiac tissue dissection, and Seahorse Metabolic Analysis. For rodent studies, the heart must be properly flushed after extraction to arrest the heart for tissue slicing. Proficiency in cardiac anatomy and experience with a vibrating microtome are essential for generating uniform, thin organotypic cardiac slices suitable for metabolic assessment. Precise tissue sectioning ensures consistency in oxygen diffusion, metabolic flux measurements, and well-to-well reproducibility.

Expertise in Seahorse Metabolic Analysis is necessary to optimize the assay for tissue-based studies. This includes a strong understanding of oxygen consumption rate (OCR) and extracellular acidification rate (ECAR) measurements, as well as the ability to modify assay conditions, media composition, and inhibitor concentrations to meet tissue-specific metabolic demands.

### Limitations of the current protocol

While assessing mitochondrial respiration, organotypic cardiac slices provide a more physiologically relevant model compared to isolated cell studies, this approach has certain limitations. Notably, electrical and mechanical stimulation of cardiac slices is not incorporated during Seahorse Metabolic Analysis, which may impact excitation-contraction-associated metabolic activity. Additionally, mechanical damage can occur during tissue sectioning, potentially affecting cell viability and metabolic measurements. Like all plate-based assays, this method is subject to plate-to-plate variability, which must be accounted for through proper experimental design and normalization strategies.

## MATERIALS

### Biological Materials

We have successfully performed the Seahorse Mito Fuel Flex Test on left ventricular tissue collected from C57BL/6J mice (JAX Laboratory, 000664) and donor human hearts that are not accepted for transplant. Other species can also be used.

#### Caution

All experiments must adhere to institutional and government guidelines. All animal experiments were approved by the Institutional Animal Care and Use Committee at Northwestern University and George Washington University and were performed in accordance with the NIH Guide for the Care and Use of Laboratory Animals. Deidentified human donor hearts that were not used in transplantation were deemed Institutional Review Board (IRB) exempt by Northwestern and George Washington Universities.

### Reagents

- Etomoxir (Cayman Chemical, cat. no. 11969)
- BPTES (Cayman Chemical, cat. no. 19284)
- UK5099 (Cayman Chemical, cat. no. 16980)
- BSA-Palmitate Saturated Fatty Acid Complex (Cayman Chemical, cat. no. 29558)
- Cell-Tak™ (Corning®, cat. no. 354240)
- Seahorse XF Calibrant Solution (Agilent Seahorse XF, cat. no. 100840-000)
- Seahorse XF DMEM medium (Agilent Seahorse XF, cat. no. 103575-100)
- Seahorse XF 1.0 M glucose solution (Agilent Seahorse XF, cat. no. 103577-100)
- Seahorse XF 100 mM pyruvate solution (Agilent Seahorse XF, cat. no. 103578-100)
- Seahorse XF 200 mM glutamine solution (Agilent Seahorse XF, cat. no. 103579-100)
- Seahorse XFe96 FluxPaks (Agilent Seahorse XF, cat. no. 102601-100)
- Sodium bicarbonate (Millipore Sigma, cat. no. S5761)
- Dulbecco′s Phosphate Buffered Saline (Millipore Sigma, cat. no. D8662)
- Dimethyl sulfoxide (Millipore Sigma, cat. no. D8418)
- Sodium Chloride (Sigma Aldrich, cat. no. S9625-1KG)
- Potassium Chloride (Sigma Aldrich, cat. no. P9333-1KG)
- Magnesium Chloride Hexahydrate (Sigma Aldrich, cat. no. M9272-1KG)
- Calcium Chloride (Sigma-Aldrich, cat. no. C1016)
- Sodium Bicarbonate (Sigma-Aldrich, cat. no. S6014-1KG)
- UltraPure™ Low Melting Point Agarose (Thermofisher, cat. no. 16520100)
- Deionized water
- 100% oxygen tank
- TissueSeal (GoBioMed, cat. no. TS1050044FP)
- AICAR (Abcam, cat. no. ab120358)
- Blebbistatin (Cayman chemicals, cat. no. 13186)
- Collagenase II (Worthington Biochemical, cat. no. LS004176)
- Collagenase IV (Thermo Fisher Scientific, cat. No. 17104019)
- Proteinase XXIV (Millipore Sigma, cat. No. P8038-100MG)
- Creatine (Sigma-Aldrich, cat. no. C3630-100G)
- Ethylenediaminetetraacetic Acid (Sigma-Aldrich, cat. no. EDS-100G)
- Fetal Bovine Serum (Thermo Fisher Scientific, cat.no. A5670701)
- Taurine (Millipore Sigma, cat. no. T8691-25G)
- HEPES (Sigma-Aldrich, cat. no. H3375-1KG)
- 2,3-Butanedione monoxime (Sigma-Aldrich, cat. no. B0753-1KG)
- Sodium Phosphate Monobasic (Sigma-Aldrich, cat. no. S0751-500G)
- Gibco™ DMEM, high glucose (Fisher Scientific, cat. no. 11-965-092)
- Insulin-Transferrin-Selenium-Ethanolamine (ITS -X) (100X) (Thermofisher, cat. no. 51500056)
- Antibiotic-Antimycotic (100X) (Thermofisher, cat. no. 15240112)
- Isoflurane, 5% (vol/vol) (Covetrus, cat. no. 029405)

**Caution:** Isofluorane is harmful to inhale, digest, or come into contact with skin. It is recommended to wear gloves and protective clothing to minimize skin contact.

### Equipment

- 2 mm biopsy punch (MedBlades)
- Single channel 20 μl micropipette (Eppendorf)
- Single channel 200 μl micropipette (Eppendorf)
- Single channel 1000 μl micropipette (Eppendorf)
- Multichannel 300 μl micropipette (Eppendorf)
- 37°C non-CO_2_ incubator (ThermoFisher)
- 37°C CO_2_ Incubator (ThermoFisher)
- Fixed Speed Vortex Mixer (Ohaus)
- Evos M5000 (ThermoFisher)
- Vibrating microtome (Campden Instruments)
- pH meter (Hanna Instruments)
- Corning® cell strainer (Millipore Sigma, cat. no. CLS431752)
- Dritz Home 3/4” White Plastic Drapery Rings 24pc (Joann, cat. no. 10702181)
- 35 mm petri dish (ThermoFisher, cat. no. 121V)
- Corning™ Centrifuge Tubes with CentriStar™ Cap (Fisher Scientific, cat. no. 05-538-59B)
- Dumont Tweezers #5 Bent 45° (WPI, cat. nom. 14101)
- Seahorse XFe96 Analyzer (Agilent Technologies)
- Centrifuge (Thermo Sorvall ST 8)

### Software

- Seahorse Wave Desktop Software (Agilent Technologies)
- ImageJ (National Health Institute)

### Reagent Setup

- **Seahorse Metabolic Analysis Cartridge:** The day before the assay (Day −1), hydrate the cartridge by adding 200 μL of Seahorse XF Calibrant to each well and leave the cartridge overnight in a non-CO_2_ incubator at 37⁰C.
- **Agarose Preparation for Organotypic Slice Slicing:** Prepare 4% agarose by dissolving 4 g of agarose in 100 mL of DI water and store at 4⁰C.
- **Solution Preparation for Cardiac Organotypic Slice Collection:** Prepare cardioplegia for heart extraction and slicing. Cardioplegia is composed of 110 mM NaCl, 16 mM KCl, 16 mM MgCl_2_, 1.5 mM CaCl_2_, and 10 mM NaHCO_3_, pH at 7.4. This should be stored at 4⁰C.
- **Seahorse Microculture Plate Coating:** Dilute 3.75 μL of Corning® Cell-Tak™ Cell and Tissue Adhesive in 6.25 μL of deionized water (DI) for each of the 96 wells that will be used in the assay, to obtain a surface coverage area of 5 μg/cm^2^. Add 10 μL of the diluted Cell-Tak™ to each well in the microculture plate. Add 30 μL of 0.1 M sodium bicarbonate to each well for a minimum of 20 minutes. Remove excess liquid. The Cell-Tak™ coated microwell plate can be stored for up to 1 week at 2-8⁰C.
- **IC_50_ Inhibitor Preparation:** Reconstitute etomoxir in dimethyl sulfoxide (DMSO) to stock concentrations of 4 mM, 8 mM, and 12.8 mM. Reconstitute BPTES in DMSO to 7.68 mM, 8.65 mM, and 9.6 mM. Reconstitute UK5099 in DMSO to 2 mM, 4 mM, 8 mM, and 12.8 mM. On the day of the assay (Day 0), prepare working dilutions of each inhibitor for the Mito Fuel Flex Test. Calculate the number of wells, fuel pathway tests, and the corresponding volume required for each cartridge injection port. Each inhibitor will be diluted 10-fold upon injection into the assay media during the assay.
- **FA Media Optimization:** On the day of the assay (Day 0), prepare BSA–palmitate supplemented Seahorse media at final concentrations of 0.0545 mM, 0.161 mM, and 0.484 mM BSA–palmitate in standard Seahorse assay media. To prepare standard Seahorse media, combine 9.7 mL Seahorse XF DMEM (pH 7.4) with 100 µL XF 1.0 M glucose, 100 µL XF 100 mM pyruvate, and 100 µL XF 200 mM glutamine. Calculate the total volume of media required based on the number of wells, using 180 μL per well.
- **Optimized Seahorse BSA-Palmitate Media:** On the day of the assay (Day 0), prepare Seahorse media supplemented with BSA-palmitate. Each well requires 180 μL of Seahorse media. To make 10 mL of media, combine 9.7 mL of Seahorse XF DMEM media, 100 µL of Seahorse XF 1.0 M glucose solution, 100 µL of Seahorse XF 100 mM pyruvate solution, 100 µL of Seahorse XF 200 mM glutamine solution, and 0.161 mM BSA-palmitate.
- **Optimized Seahorse Mito Fuel Flex Test Inhibitors for Cardiac Organotypic Slices:** Reconstitute etomoxir, BPTES, and UK5099 in DMSO to a stock concentration of 8 mM, 9.6 mM, and 8.0 mM, respectively. On the day of the assay (Day 0), dilute stock inhibitors for the Mito Fuel Flex Test Assay and calculate the number of wells, number of fuel tests, and volume required for each cartridge port. The final well concentration for each inhibitor for all fuel tests should be 0.80 mM, 0.96 mM, and 0.8 mM for etomoxir, BPTES, and UK5099, respectively.
- **Cell Isolation Buffer Preparation:** On the day of the experiment for primary adult cell isolation, prepare the basal buffer containing 130 mM NaCl, 5 mM KCl, 0.5 mM NaH₂PO₄, 10 mM HEPES, 10 mM glucose, 10 mM 2,3-Butanedione monoxime (BDM), and 10 mM taurine. This basal buffer serves as the foundation for other solutions. The Ethylenediaminetetraacetic Acid (EDTA) buffer is prepared by adding EDTA to the basal buffer to achieve a final concentration of 5 mM. The perfusion buffer is made by supplementing the basal buffer with MgCl₂ to reach a final concentration of 1 mM. The digestion buffer is prepared by adding protease XIV at 0.05 mg/mL, collagenase II at 0.5 mg/mL, and collagenase IV at 0.5 mg/mL to the basal buffer. The digestion buffer needs to be pre-warmed to 37 ⁰C. The stop buffer is made with perfusion buffer by adding 5% fresh Fetal Bovine Serum (FBS) on the day of the experiment. The stop buffer is kept at room temperature.

### Equipment Setup

- **Vibrating Microtome Calibration:** Load a ceramic blade into the blade holder and attach the calibration unit provided with the microtome. Adjust the calibration screws until the z-axis alignment is below 1 μm. Set the cutting thickness to 100 μm, the advance speed to 0.04 mm/s, the horizontal vibrational amplitude to 2 mm, and the vibration frequency to 80 Hz. The calibration can be completed before Day 0. On the day of the assay (Day 0), fill the vibrating microtome bath with 4⁰C cardioplegia solution, and keep the bath chilled by filling the bath’s surroundings with ice (replace ice as needed). Oxygenate the solution by bubbling 100% O_2_ throughout the experiment. Set up a recovery dish next and fill with cardioplegia solution at room temperature and bubble with O_2_ throughout the experiment. Fill the recovery dish with cell strainers and washers as needed.

## PROCEDURE

### Tissue Collection Methods

A. **Mouse Left Ventricle (LV) Free Wall Dissection (**15 minutes per animal)

**1.** Anesthetize the mouse using 5% isoflurane with 1 mL/min oxygen flow in an induction chamber for several minutes until the mouse becomes unconscious. Confirm loss of consciousness using the toe-pinch method.
**2.** Perform cervical dislocation, followed by a thoracotomy.
**3.** Extract the heart and cannulate the aorta.
**4.** Retrogradely perfuse the heart with 4⁰C cardioplegia.
**5.** Isolate the LV-free wall and temporarily store (up to 6 hours) the tissue in 4⁰C cardioplegia.
B. **Human Donor Heart Dissection** (**10 min**)

**6.** Dissect the LV free wall, right ventricular (RV) free wall, RV outflow tract (RVOT), right atrium (RA), and left atrium (LA) from a donor human heart.

**Organotypic Slice Collection Positioning** (1 hour per heart)

**7.** Glue an agarose gel to the bottom and back of the metal tissue holders of the vibrating microtome with TissueSeal. Adhere the tissue to the agarose gel, ensuring firm contact between both the bottom and back agarose blocks.
**8.** For the mouse LV, position the free wall with the endocardium facing down; for human donor heart tissue, position the epicardium facing down. Gently adhere the tissue to the agarose gel, ensuring firm contact with both the base and back of the agarose blocks.

**Critical Step:** Ensure that the tissue is flat and properly supported by the vertical agarose gel. Avoid applying excessive tissue glue, as this may cause the tissue to become coated in glue, which can interfere with proper adhesion and tissue oxygenation during sectioning.

**9.** Transfer the metal tissue holder to the vibrating microtome bath, ensuring the tissue is fully submerged in ice-cold cardioplegia bubbled with O₂.
**10.** Position the blade above the tissue and raise the bath until the blade is just above the tissue surface. This sets the appropriate height to begin sectioning.

**Critical Step:** Adjust the initial cutting height carefully to optimize tissue preservation and efficiency. If the blade is too close to the tissue, excess tissue may be lost before proper slicing begins, potentially reducing the number of usable slices. On the other hand, if the blade is too far, the cutting process will be inefficient, requiring additional time to reach the desired tissue depth.

**11.** Position the blade at the front edge of the tissue and initiate slicing.
**12.** Once a slice is generated, collect it using a cell strainer with an agarose base and a mesh top to maintain a flat orientation. Transfer the slice to the recovery bath containing room-temperature cardioplegia bubbled with O₂. Slicing can take approximately 1 hour per heart, depending on the size of the tissue and the number of replicates required for the assay. Using multiple vibrating microtomes increases throughput, allowing for simultaneous slice collection from multiple hearts, thereby improving efficiency in tissue preparation.

**Organotypic Slice Attachment and Incubation** (2 Hours)

**13.** Transfer slices from the recovery bath to a Petri dish containing Dulbecco’s Phosphate-Buffered Saline (DPBS) as needed.

**Critical Step:** Use only enough DPBS to allow the slice to lay flat. Excess solution can make biopsy punching difficult, while insufficient solution may prevent the slice from properly flattening, affecting the precision of subsequent processing.

**14.** Use a 2 mm biopsy punch to generate 2 mm mini slices, applying slight pressure to create clean and precise cuts.

**Critical Step**: Ensure the biopsy punch is sharp and not dulled over time, as a dull punch can make it difficult to obtain clean and precise 2 mm mini slices, potentially damaging the tissue.

**15.** Add 20 μL of DPBS to the well of the CellTak-coated microculture plate where the slice will be attached.
**16.** Transfer the 2 mm mini slice into the well.
**17.** Carefully remove the 20 μL of DPBS, ensuring the slice is flat at the bottom of the well.
**18.** Allow the slice to adhere to the microculture plate for 1 – 2 minutes before proceeding.

**Critical Step:** Allow the 2 mm mini slice to adhere for an appropriate duration. If the slice does not adhere long enough, it may detach and float when media is added. Conversely, if it adheres for too long, it may become ischemic, compromising tissue viability.

**19.** Slowly add 180 μL of warmed media to the well.

**Critical Step:** Add the media gradually to prevent the slice from detaching and floating to the top, ensuring it remains properly adhered to the well plate.

**20.** Repeat Steps 13 – 19 for the remaining wells of the assay.
**21.** Add 180 μL of Seahorse BSA-Palmitate media to four wells to serve as the background wells (negative controls).
**22.** Transfer the microculture well plate to a CO₂ incubator and incubate for 45 – 60 minutes after all 2 mm mini slices are attached.

**Loading Inhibitors into Seahorse Metabolic Analysis Cartridge Ports** (30 minutes for 90 wells)

**23.** While the microculture well plate is incubating, prepare and load inhibitors into Ports A and/or B for the Seahorse Inhibitor IC₅₀ Assay, Control Mouse Mito Fuel Flex Test, Isolated Primary Mouse Cardiomyocyte Mito Fuel Flex Test, AICAR Metabolic Flux Measurements, or Human Heart Chambers Mito Fuel Flex Test.

**Critical Step:** If a given port has to be used on the plate, ensure that all ports of that label are filled with either an inhibitor or media. For example, when Port A is used in all assay wells, all Port As on the plate have to be loaded. If some wells remain empty, the Seahorse Metabolic Instrument may fail to accurately inject the correct amount of inhibitor. For wells that will not be used in the assay, load the appropriate volume of media to maintain uniformity. Load 20 µL into Port A and 22 µL into Port B.

### Assay Overview

Multiple metabolic assays were performed to adapt Seahorse Mito Fuel Flex Test for tissue preparations. These assays include the Seahorse Inhibitor IC₅₀ Assay, Optimization of FA Media Assay, Control Mouse Mito Fuel Flex Test, Isolated Primary Mouse Cardiomyocyte Mito Fuel Flex Test, AICAR Metabolic Flux Measurements, and Human Heart Chambers Mito Fuel Flex Test.

**Half-Maximal Inhibitor Concentration (IC_50_) Assay** (2 hours)

**24.** Generate mouse LV-free wall 2 mm mini slices adhered to a Seahorse microculture plate following steps 1 – 22.
**25.** Create a Seahorse XF Custom Template where the assay is configured to include 3 baseline measurement cycles without inhibitors, followed by 6 mix and measurement cycles, each cycle lasting 3 minutes.
**26.** Pipette 20 µL of each substrate inhibitor solution into the appropriate wells of Port A in the Seahorse XFe96 sensor cartridge. The inhibitor stock concentrations tested are as follows: Etomoxir: 4 mM, 8 mM, 12 mM BPTES: 7.68 mM, 8.65 mM, 9.65 mM UK5099: 2 mM, 4 mM, 8 mM, 12.8 mM
**27.** Insert the Seahorse XFe96 sensor cartridge into the Seahorse XFe96 Analyzer and initiate the calibration process.
**28.** Once calibration is complete, remove the sensor cartridge and place the microplate into the Seahorse XFe96 Analyzer.
**29.** Initiate the pre-programmed Seahorse XF assay and proceed with data acquisition.
**30.** Proceed to Tissue Area Normalization and Analysis Section.

**Optimization of FA Media Assay** (2 hours)

**31.** Generate mouse LV-free wall 2 mm mini slices adhered to a Seahorse microculture plate following steps 1 – 18.
**32.** Slowly add 180 μL of warmed Seahorse media containing 0 mM, 0.0545 mM, 0.161 mM, or 0.484 mM BSA-palmitate and incubate for 60 min.
**33.** Create a custom Seahorse XF Template, incorporating a baseline respiration period without inhibitor injection, consisting of nine measurement cycles (6 min per cycle).
**34.** Follow Steps 28 – 30.
**35.** Proceed to the Tissue Area Normalization and Analysis Section.

**Control Mito Fuel Flex Test Assay** (2 hours)

**36.** Generate mouse LV-free wall 2 mm mini slices adhered to a Seahorse microculture plate following steps 1 – 23.
**37.** Load the Seahorse XFe96 sensor cartridge ports following the Seahorse Mito Fuel Flex Test Kit User Manual for each fuel dependency and capacity assay which will be performed. Based on the inhibitor IC_50_ test results, inhibitor concentrations of BPTES (9.6 mM stock), etomoxir (8.0 mM stock), and UK5099 (8.0 mM stock) were used for tissue (Table 1).
**38.** Create a custom Seahorse XF Template to define measurement and injection cycles. Record baseline respiration over three 3-minutes cycles, followed by Port A inhibitor injection, a mix cycle, a 0-minute wait cycle, and three additional 3-minute measurement cycles. Inject the second inhibitor, then repeat the mix cycle, a 0-minute wait cycle, and three final 3-minute measurement cycles.
**39.** Follow Steps 28 – 30.
**40.** Proceed to the Tissue Area Normalization and Analysis Section.

**Table 1.** Final Concentrations of Inhibitors For Tissue Adapted Seahorse Metabolic Analysis.

| Inhibitor | Stock Concentration (mM) | Final Well Concentration (mM) |
| --- | --- | --- |
| Etomoxir | 8.0 | 0.80 |
| UK5099 | 8.0 | 0.80 |
| BPTES | 9.6 | 0.96 |

**AICAR Metabolic Flux Measurements** (2 hours)

**41.** Generate mouse LV-free wall 2 mm mini slices adhered to a Seahorse microculture plate following steps 1 – 18.
**42.** Slowly add 180 μL of warmed Seahorse BSA-Palmitate media with either 0, 0.25, 0.5, or 1 mM AICAR to each well and incubate for 60 minutes.
**43.** Create a custom Seahorse XF Template, incorporating a baseline respiration period without inhibitor injection, consisting of nine measurement cycles (6 min per cycle).
**44.** Follow Steps 28 – 30.
**45.** Proceed to Tissue Area Normalization and Analysis Section.

**Tissue Area Normalization and Analysis** (2.5 hours for 90 wells)

**46.** Following assay completion, image each well containing a 2 mm mini slice using EVOS™ M5000 Imaging System. For 90 wells, this will take approximately 45 minutes.

**Pause Point:** After capturing images of all wells, analysis can be performed at a later time.

- Data analysis takes approximately 2 hours for 90 wells.

**47.** Calculate the area of the 2 mm mini slice in each well using ImageJ software.
**48.** Normalize the 2 mm mini slice area for each well to the largest 2 mm mini slice area on the plate.
**49.** Normalize O_2_ levels, pH levels, OCR, and ECAR based on the normalized areas.
**50.** For the IC_50_ assay, calculate the percent change by comparing the first baseline measurement and the final post-injection measurement. For the BSA-Palmitate Optimization Assay, calculate the percent change in supplemented media average OCR relative to the same mouse control media average OCR.
**51.** For the Seahorse Mito Fuel Flex Test, calculate fuel dependency (Equation 1) and capacity (Equation 2) as defined by the Seahorse Mito Fuel Flex Test Kit User Manual.

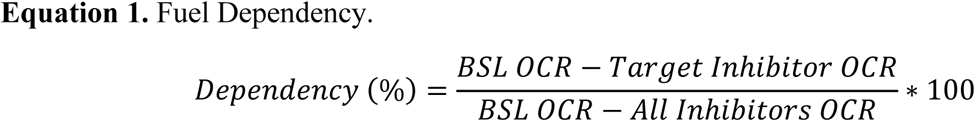

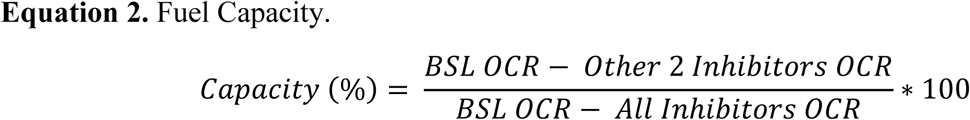

**Adult Cardiomyocyte Isolation and Culture of Adult Mouse Cardiomyocytes** (7 hours)

**52.** Excise the mouse heart as described in steps 1 – 3.
**53.** Retrogradely perfuse the heart with 9 mL EDTA buffer at a rate of 1-2 mL/min, to clear blood in vessels.
**54.** Retrogradely perfuse the heart with 3 – 4 mL perfusion buffer. This was used to perfuse the heart to wash out the EDTA buffer.
**55.** Connect the heart to a Langendorff Perfusion system, and retrogradely perfuse the heart with 37⁰C digestion buffer beginning at a flow rate of 2 mL/min and increasing the flow rate to 3 – 4 mL/min as digestion processes.

**Critical Step:** Digestion should take 10 – 15 minutes, depending on the age of the heart. Prolonged digestion can lead to warm ischemia and cell damage, compromising cell yield and viability.

**56.** Remove the heart from the Langendorff System and isolate the LV in a 35 mm petri dish filled with 2 mL of fresh digestion buffer.
**57.** Gently tease apart the LV into small pieces using fine-tip forceps.
**58.** Gently pipette to fully dissociate the tissue using transfer pipettes.
**59.** Add 2 mL room-temperature stop buffer to quench enzymatic activity.
**60.** Filter cell-tissue suspension into a 50 mL or a 15 mL conical tube using a 200 μm cell strainer to remove tissue debris.
**61.** Wash the strainer and petri dish with the stop buffer to collect the remaining cells.
**62.** Centrifuge the conical tube at 100 g, for 2 min, at room temperature to pellet rod-shaped cardiomyocytes.
**63.** Add the stop buffer to resuspend cells. The volume of stop buffer added depends on the pellet size.
**64.** Count cells by hemocytometer.
**65.** Remove laminin from each well and gently wash with DPBS once.
**66.** Reintroduce calcium stepwise to avoid the calcium paradox. Add 100 mM of CaCl_2_ every 3 minutes to restore Ca^2+^ concentrations to 0.2 mM, 0.4 mM, 0.6 mM, 1mM, 1.4 mM, and 1.8 mM.
**67.** Centrifuge conical tube at 100 g, for 2 min, at room temperature to pellet rod-shaped cardiomyocytes.
**68.** Add warmed culture media (DMEM-HG, 5% FBS, 1% ITS, 1% Anti-anti, 10 μM Blebbistatin) to resuspend cells.
**69.** Dispense cell suspension into the Seahorse 96-well microculture plate, ensuring each well has at least 500 cardiomyocytes.
**70.** Allow cardiomyocytes to adhere for at least 3 hours in an incubator.
**71.** Aspirate culture media (DMEM-HG, 5% FBS, 1% ITS, 1% Anti-anti, 10 μM Blebbistatin).
**72.** Add 180 μL of Seahorse BSA-palmitate media and incubate for 45 – 60 minutes.
**73.** Perform Seahorse Mito Fuel Flex Test following inhibitor concentrations as provided in User Manual for BPTES, etomoxir, and UK5099.

**Adult Cardiomyocyte Normalization and Analysis** (30 minutes)

**74.** Normalize each well based on the number of cells dispensed into that well.
**75.** Normalize O_2_ levels, pH levels, OCR, and ECAR based on the normalized cell number.
**76.** Repeat Step 47.

### Timing

- Steps 1 – 5: 15 min per animal
- Step 6: 10 min
- Steps 7 – 11: 5 min per heart
- Steps 12: 1 hour per heart (Note this may vary depending on tissue size)
- Step 13 – 21: 1 hour for 90 wells
- Step 22: 45 minutes to 1 hour
- Steps 23: 30 minutes for 90 wells
- Steps 24 – 30: 2 hours
- Steps 31 – 35: 2 hours
- Steps 36 – 40: 2 hours
- Steps 41 – 45: 2 hours
- Step 46: 45 minutes for 90 wells
- Steps 47 – 51: 2 hours for 90 wells
- Steps 52 – 73: 7 hours
- Steps 74 – 76: 30 min.

### Troubleshooting

If there is no change in OCR after inhibitor injections, it may indicate that the tissue slice detached from the well plate or that the system failed to inject inhibitors properly. Detached slices can lead to inaccurate metabolic measurements. To prevent detachment, ensure slices are given adequate time to adhere before adding media and inhibitors. If the issue is due to incomplete inhibitor injection, verify that all injection ports released the inhibitors after the assay is complete. Wells showing these abnormalities should be excluded from the final analysis.

## ANTICIPATED RESULTS

To establish optimal assay conditions, inhibitor concentrations and Seahorse media composition were optimized for cardiac tissue. These optimizations enabled the successful execution of the Mito Fuel Flex Test on healthy living mouse LV organotypic cardiac slices (Figure 1A). For comparison, the standard cell-based Seahorse Mito Fuel Flex Test was performed on isolated primary adult mouse cardiomyocytes (Figure 1B). Additionally, Seahorse Metabolic Analysis dose-dependently detected both increases and decreases in baseline respiration, confirming its sensitivity in measuring metabolic changes. Lastly, this methodology was applied to human cardiac organotypic slices, collected from the left and right atria (LA and RA), left and right ventricles (LV and RV), and right ventricular outflow tracts (RVOT), allowing for the assessment of chamber-specific metabolic profiles of the human heart.

**Figure 1.**
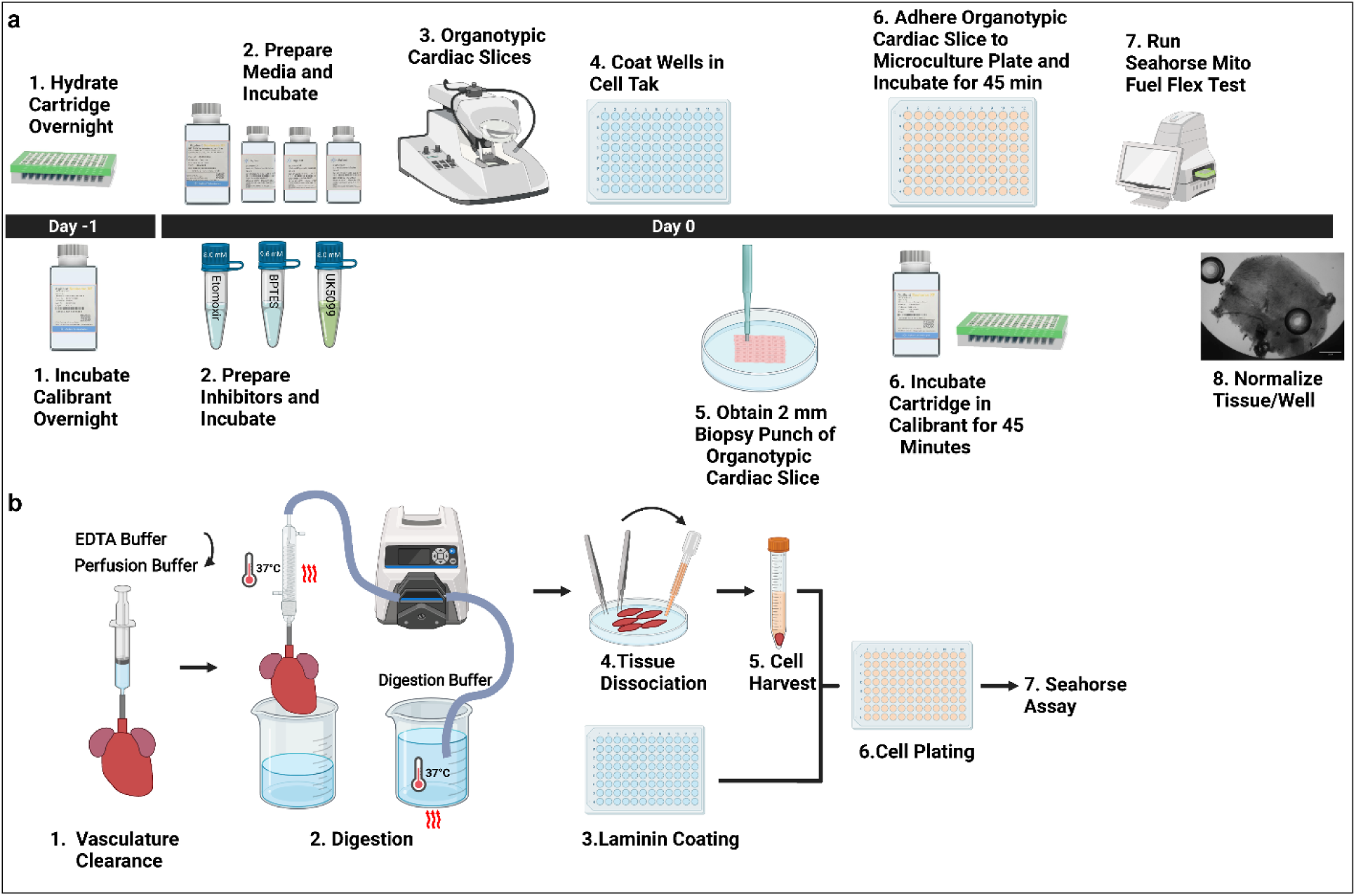
Schematic of Protocol for Seahorse Metabolic Analysis Performed with Cardiac Organotypic Slices and Isolated Primary Cardiomyocytes. **A)** The protocol starts with cartridge hydration and calibrant incubation overnight. On the assay day, media, inhibitors, and cardiac slices are prepared. Microculture wells are coated with CellTak, and 2 mm biopsy punches of slices are attached. After approximately 45 minutes of incubation, the Seahorse Assay is performed. Recorded OCR and ECAR values are normalized to the tissue area in each well. **B)** The isolation protocol starts with retrograde perfusion to clear the blood and calcium out of vessels, followed by digestion through a simplified Langendorff system. Isolated cells are plated on pre-coated plates for Seahorse assays. Created in BioRender. Amaral, P. (2025) https://BioRender.com/5mvch9x

### Optimization of Inhibitor Concentrations for Seahorse Mito Fuel Flex Test

To determine IC_50_ for the Seahorse Mito Fuel Flex Test, an inhibitor dose-response study was conducted using mouse LV organotypic cardiac slices. Representative O_2_ levels and pH traces at baseline and after inhibitor treatment demonstrate a reduction in the rate of oxygen consumption by the tissue and an increase in the rate of change of pH in the media (Figures 2A and 2B). Following inhibitor treatment, a decrease in OCR and a concurrent increase in ECAR were observed (Figures 2C and 2D). The inhibitor stock concentrations required to achieve at least a 50% reduction in OCR were determined to be 9.6 mM for BPTES, 8.0 mM for etomoxir, and 8.0 mM for UK5099 (Figure 2E). These concentrations also resulted in an increase in ECAR for BPTES, etomoxir, and UK5099 (Figure 2F). A summary of optimized stock and final well concentrations for BPTES, etomoxir, and UK5099 is provided in Table 1.

**Figure 2.**
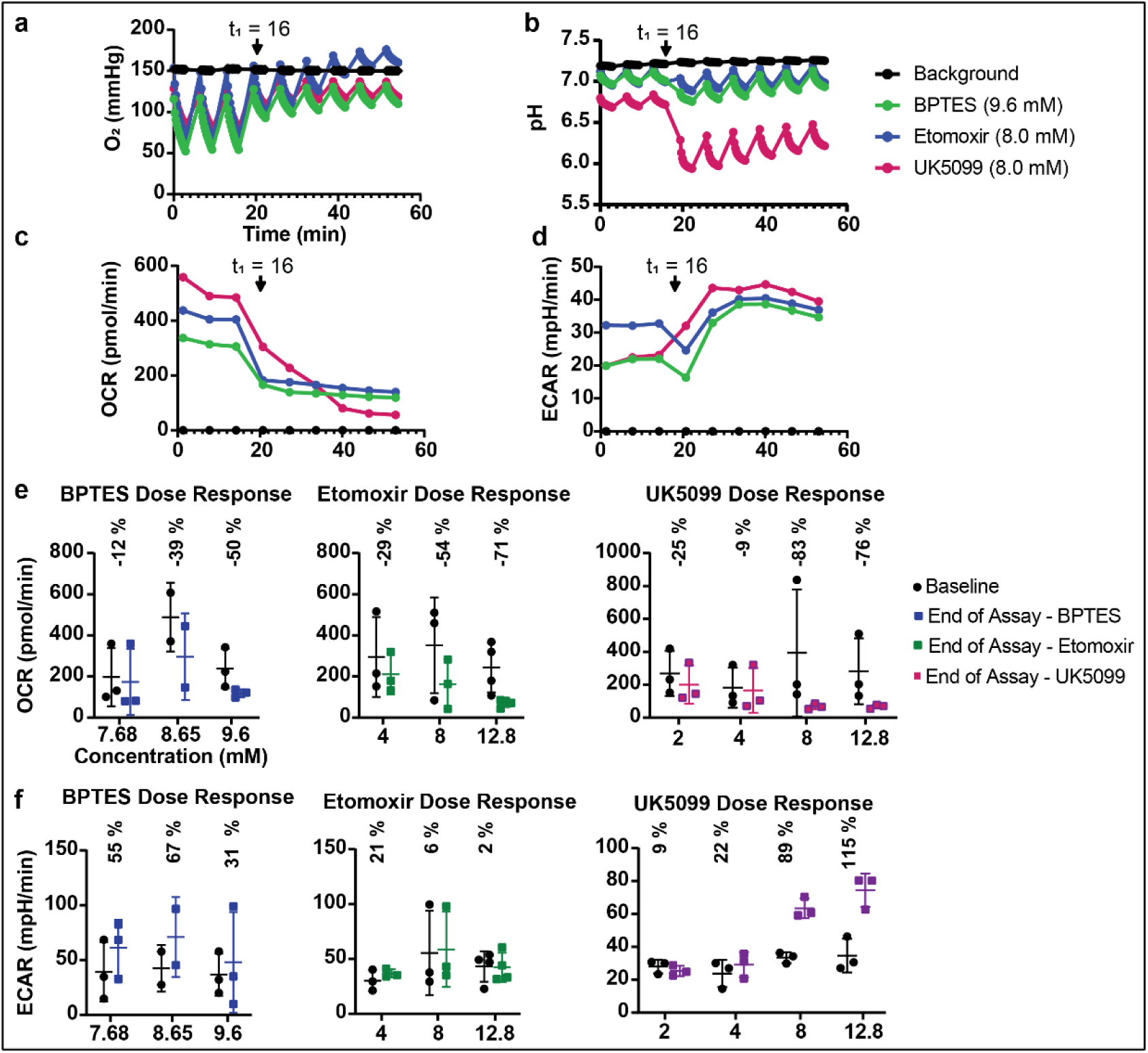
Seahorse Metabolic Analysis Mito Fuel Flex Test Inhibitor Dose Response. **A, B)** Representative oxygen consumption levels (O_2_) and extracellular acidification levels (pH). The arrow at t_1_ indicates the time of inhibitor injection. **C, D)** Representative oxygen consumption rate (OCR) and extracellular acidification rate (ECAR) dose-response were calculated from changes in O_2_ and pH levels. **E, F)** OCR and ECAR at baseline (before inhibitor injection) and at the end of the assay are shown for BPTES (blue), etomoxir (green), and UK5099 (magenta) at varying concentrations. The percentage change between the baseline value and the end of the assay is listed above each group.

### Optimized BSA-palmitate supplemented Seahorse Media

The Seahorse Mito Fuel Flex Test User Manual recommends a standard Seahorse media formula composed of DMEM culture media, glucose, pyruvate, and glutamine. However, FA, the primary energy fuel of a healthy heart, is notably absent from this media composition. To address this, Seahorse media was supplemented with BSA-palmitate at concentrations of 0 mM, 0.0545 mM, 0.161 mM, and 0.484 mM to evaluate metabolic response. Representative O_2_ level, pH, OCR, and ECAR traces are illustrated for each Seahorse culture media condition. (Figure 3A – 3D). An energy map, which plots glycolytic activity (ECAR) on the x-axis and mitochondrial respiration (OCR) on the y-axis, illustrates the group responses to media supplemented with BSA-palmitate (Figure 3E). Average OCR and ECAR measurements per condition per mouse (Figures 3F and 3H) and the percent change relative to control Seahorse media (Figures 3G and 3I) were calculated for each BSA-palmitate supplemented Seahorse media concentration. We selected the BSA-palmitate supplemented media concentration that had the maximum change in OCR percent change relative to control media, which was 0.161 mM. This optimized media composition more accurately replicates the metabolic environment of cardiac tissue, improving the reliability of Seahorse Metabolic Analysis in cardiac organotypic slices.

**Figure 3.**
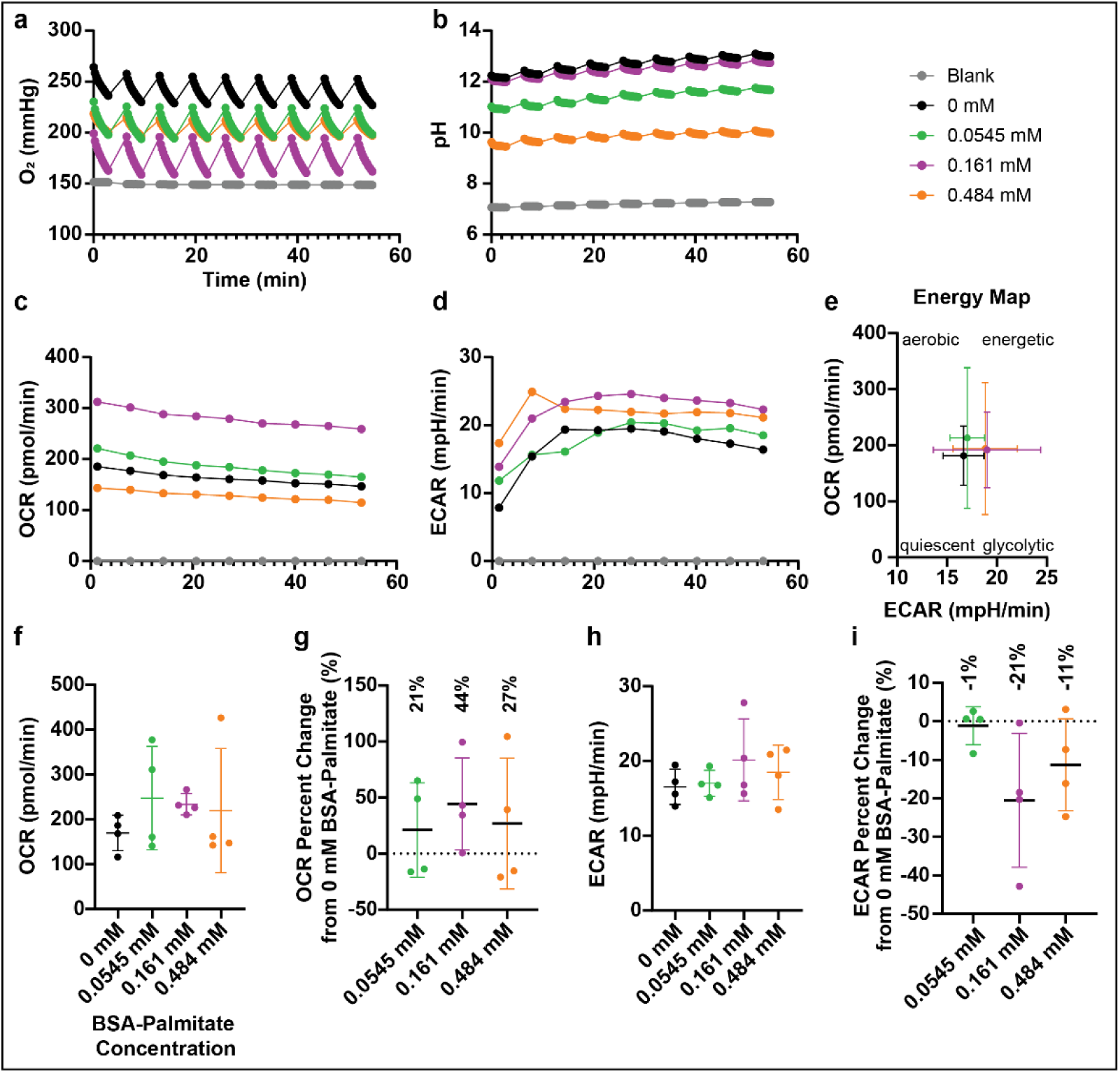
BSA-palmitate supplemented Seahorse media increases OCR. **A, B)** Representative O_2_ and pH levels were recorded using control Seahorse culture media (0 mM) and media supplemented with 0.0545 mM, 0.161 mM, and 0.484 mM BSA-palmitate. **C, D)** Representative OCR and ECAR curves were derived from changes in O_2_ and pH levels across the media conditions described. **E)** Energy map plotting baseline ECAR and OCR values using culture media described. **F)** Average OCR per mouse at varying BSA-palmitate concentrations. **G)** Percent change in OCR relative to control media at varying BSA-palmitate concentrations for each mouse. The maximum percent change in OCR (44%) was observed with 0.161 mM BSA-palmitate. **H)** Average ECAR per mouse at varying BSA-palmitate concentrations. **I)** Percent change in ECAR relative to control media at varying BSA-palmitate concentrations.

### Healthy Mouse Seahorse Mito Fuel Flex Test

Using the optimized Seahorse Mito Fuel Flex Test inhibitor concentrations and Seahorse media, the assay was performed on healthy mouse LV organotypic slices as described in the procedure. Representative O_2_ and pH levels measured, and the calculated OCR and ECAR values are shown for each of the 6 tests in the assay (Figure 4A – 4D). The energy map in Figure 4E illustrates the individual energy profile of each mouse and the average energy profile of all mice (black). Dependency on FA, GLC, and GLN was calculated (Figure 4F). No significant differences in metabolic dependency on any given substrate were measured. This indicates that in control conditions, when the heart tissue is not stressed and is cultured under static conditions as described above, metabolism is equally dependent on all three substrates – FA, GLC, and GLN. Finally, the metabolic capacity of each individual substrate was measured (Figure 4G). Metabolic capacity for GLN was significantly lower than FA metabolism under these culture conditions. This metabolic profile reflects the expected substrate utilization in cardiac tissue, highlighting the roles of FA and GLC as the primary energy sources under static ex vivo conditions.

**Figure 4.**
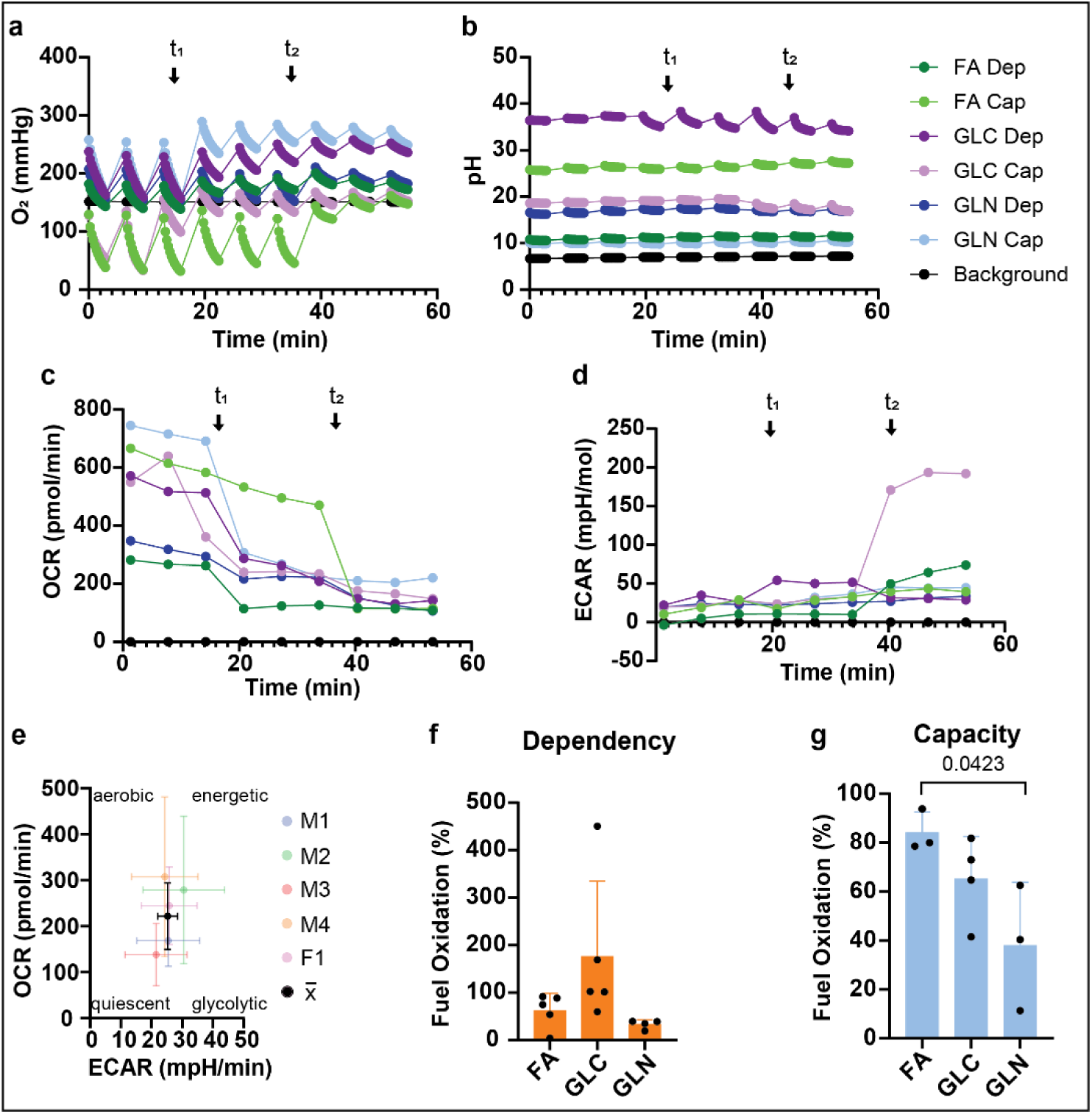
Seahorse Mito Fuel Flex Test performed on mouse organotypic cardiac slices. **A, B)** Average oxygen (O_2_) and pH level traces measured for background, representative FA Dep, FA Cap, GLC Dep, GLC Cap, GLN Dep, and GLN Cap wells are shown. First and second inhibitor port injections are indicated at time points 1 (t_1_) and 2 (t_2_). **C, D)** Representative OCR and ECAR were calculated for the groups mentioned above from changes in O_2_ and pH levels. **E)** Energy map plotting average baseline ECAR and OCR for each mouse (displayed transparently, n = 5) and the collective average of all mice baseline ECAR/OCR (displayed in black, x̅). **F, G)** Dependency and capacity were calculated for each mouse. A one-way ANOVA with multiple comparisons was performed to determine statistical significance.

### Isolated Primary Adult Cardiomyocyte Seahorse Mito Fuel Flex Test

The Seahorse Mito Fuel Flex Test was performed on isolated primary adult mouse cardiomyocytes following the Agilent User Manual inhibitor concentrations and with our optimized Seahorse media to compare differences in fuel utilization in cells. Representative O_2_ and pH levels, along with the calculated OCR and ECAR, are shown for each fuel test (Figures 5A – 5D). An energy plot displaying individual metabolic profiles for each mouse from which cardiomyocytes were isolated, and the average baseline respiration for all mice is shown in Figure 5E. No significant differences in metabolic dependency or capacity for fatty acids (FA), glucose (GLC), or glutamine (GLN) were observed, suggesting that under basal, unstressed conditions, isolated primary adult cardiomyocytes utilize all three substrates equally (Figures 5F and 5G). These findings highlight differences between isolated cardiomyocytes and intact cardiac tissue, emphasizing the impact of cell isolation and culture conditions on metabolic preference. Isolated cardiomyocytes undergo shifts in metabolism, favoring anaerobic glycolysis over aerobic oxidative phosphorylation^26^. Additionally, the isolation process of primary adult cardiomyocytes involves proteolytic enzymes that remove extracellular glycoproteins that regulate metabolic functions^26^. These factors could contribute to the metabolic substrate capacity observed between cardiac slices and isolated cardiomyocytes.

**Figure 5.**
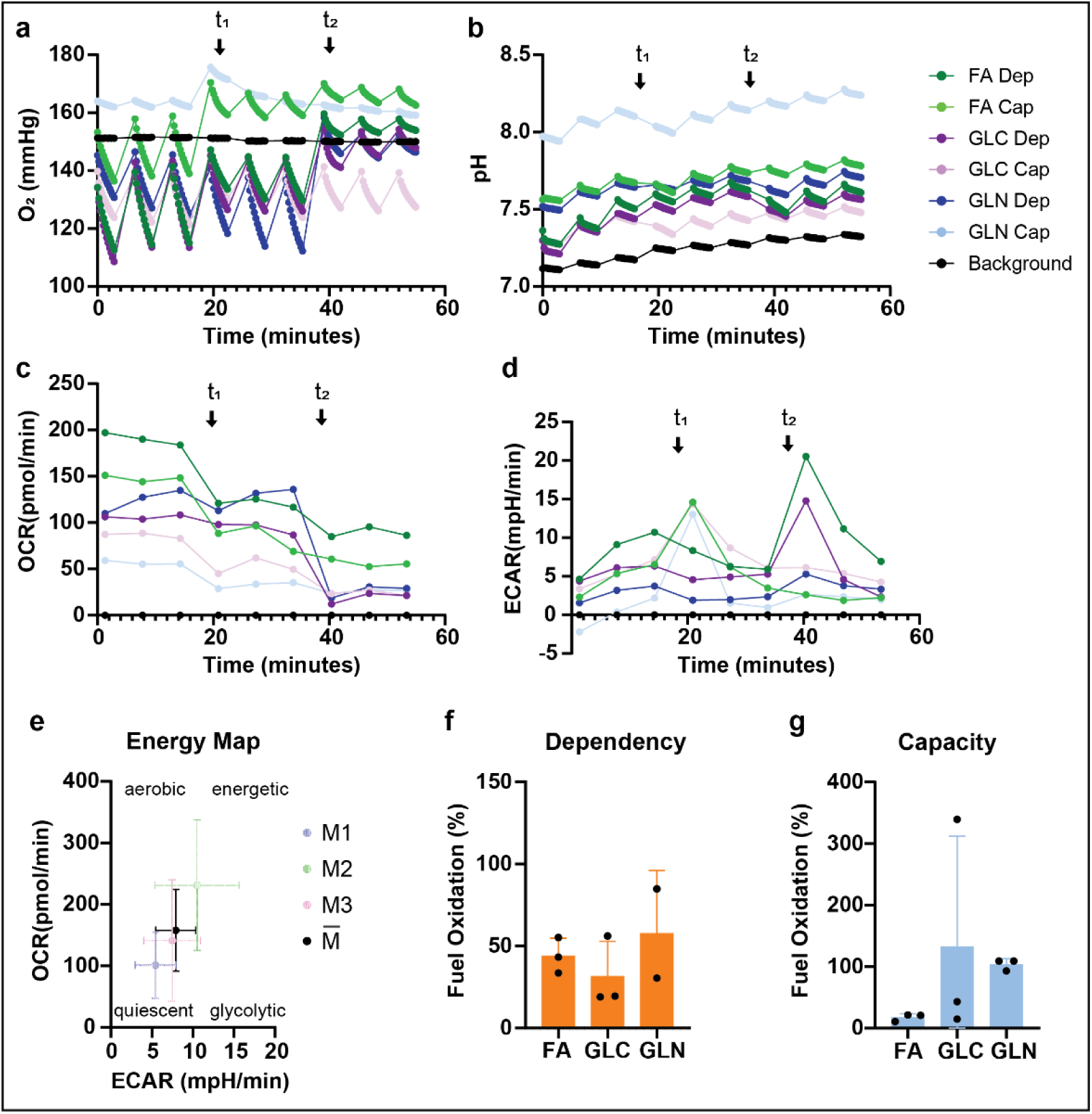
Seahorse Mito Fuel Flex Test performed on isolated primary mouse cardiomyocytes. **A, B)** Representative O_2_ and pH levels recorded isolated primary mouse cardiomyocytes during the Seahorse Mito Fuel Flex Test. First and second inhibitor port injections are indicated at time points 1 (t_1_) and 2 (t_2_). **C, D)** Representative OCR and ECAR curves calculated from changes in O_2_ and pH levels. **E)** Energy map plotting average baseline ECAR and OCR for each mouse (displayed transparently, n = 3) and the collective average of all mice baseline ECAR/OCR (displayed in black, x̅). **F, G)** The dependency and capacity for each fuel were calculated for each mouse, respectively. A one-way ANOVA with multiple comparisons was performed to determine statistical significance.

### AICAR Results

AICAR is an AMPK activator known to enhance metabolism, but at high concentrations, it has been reported to suppress metabolic activity^27–29^. Next, seahorse analysis was performed on mouse LV slices treated with varying doses of AICAR – 0 mM (Control media), 0.25 mM, 0.5 mM, and 1 mM. BSA-palmitate-supplemented media were used as described above.

Representative O_2_ and pH levels, along with the calculated OCR and ECAR are shown for each fuel test (Figures 6A – 6D). An energy map illustrates the dose-dependent metabolic enhancement, with slices treated with 0.25 mM and 0.5 mM AICAR indicating an increased energetic profile (shift to upper right quadrant), while 1 mM AICAR treatment resulted in a significant metabolic decline (Figure 6E, shift to bottom left quadrant). Specifically, 1 mM AICAR treatment led to a decrease in OCR (p = 0.05) compared to 0.5 mM AICAR (Figure 6F). Similarly, ECAR trended downward with 1 mM AICAR treatment (p = 0.08), suggesting a potential toxic effect at high concentrations (Figure 6G).

**Figure 6.**
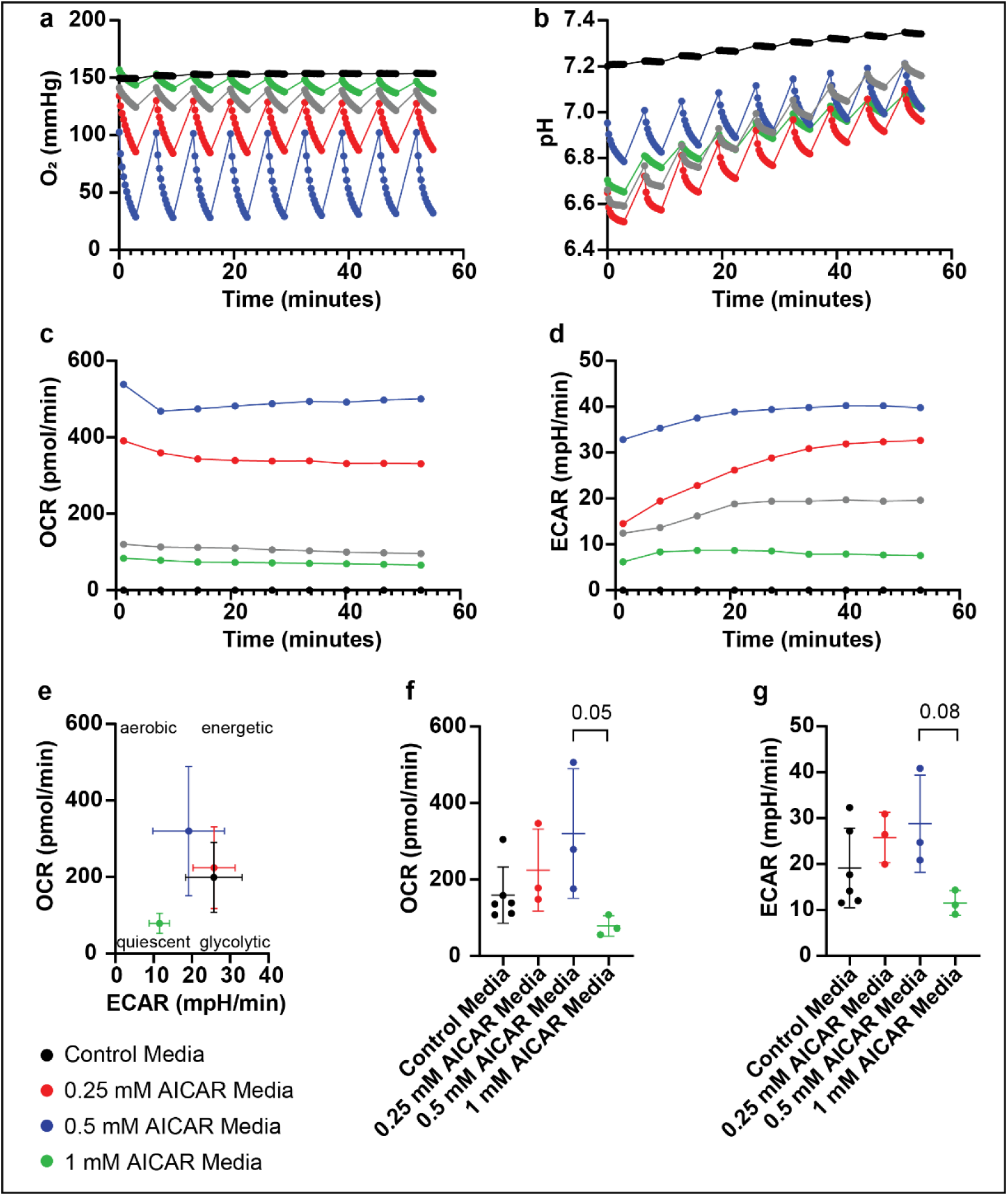
Seahorse Mouse Mito Fuel Flex Test AICAR Dose Response. **A, B)** Representative O_2_ and pH levels were recorded with BSA-palmitate supplemented Seahorse Culture Media treated with 0, 0.25 mM, 0.5 mM, and 1 mM AICAR. **C, D)** Representative OCR and ECAR curves were calculated from changes in O_2_ and pH levels in the groups described above. **E)** Energy map plotting ECAR and OCR values of the culture media described. **F, G)** The average OCR and ECAR at the end of the assay for each culture condition for each mouse is shown, where there is a trend toward increased metabolism with increasing AICAR concentrations for 0.25 mM and 0.5 mM AICAR, while 1 mM AICAR was toxic. A one-way ANOVA with multiple comparisons was performed to determine statistical significance.

**Figure 6.**
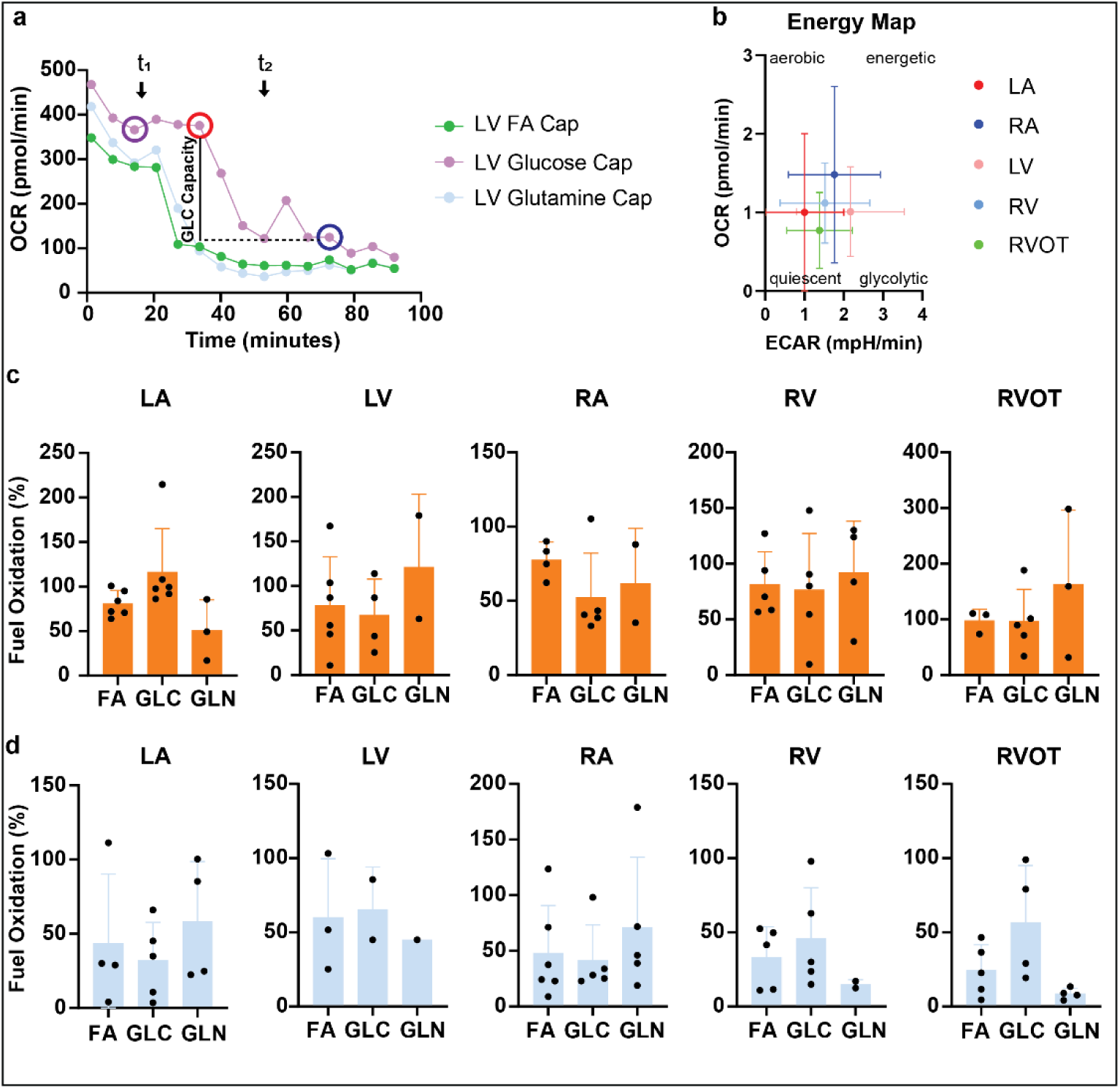
Human LV demonstrates increased glycolytic activity compared to the LA and RVOT. **A)** Representative OCR curve demonstrating GLC capacity measurement definition. Circles indicate the specific OCR measurements used in the fuel capacity analysis: baseline OCR (purple), OCR following inhibition of alternative fuel pathways (red), and OCR following inhibition of the target fuel pathway (blue). These values were used to calculate fuel capacity, as defined in Equation 2. **B)** Energy map displaying the BSL ECAR and OCR values for each chamber and RVOT. **C)** Fuel dependency plotted for each chamber and RVOT. **D)** Fuel capacity plotted for each chamber and RVOT. A one-way ANOVA with multiple comparisons was performed to determine statistical significance.

### Chamber-Specific Metabolic Differences in Human Organotypic Cardiac Slices

The optimized Seahorse Mito Fuel Flex Test protocol was applied to human left atrial (LA), LV free wall, right atrial (RA), right ventricular (RV) free wall, and right ventricular outflow tract (RVOT) organotypic slices to assess chamber-specific metabolic profiles. The assay utilized inhibitor concentrations optimized for organotypic cardiac slices and Seahorse media supplemented with BSA–palmitate. Representative OCR traces for FA, GLC, and GLN fuel capacities of the LV are shown (Figure 6A). The energy map illustrates metabolic differences among human heart regions. While baseline OCR did not significantly differ across regions, the LV exhibited higher baseline ECAR compared to the LA (p = 0.0383) and trended toward an increase when compared to the RVOT (p = 0.0747), suggesting a higher rate of glycolysis in the LV relative to the LA and RVOT. The increased ECAR in the LV indicates a metabolic shift toward glycolysis, which may result from the lack of mechanical and electrical stimulation during Seahorse Metabolic Analysis. This shift suggests that, under static culture conditions, the LV may rely more on glucose metabolism than in its native physiological state. FA, GLC, and GLN capacity and dependency were also compared between human heart chambers, but no significant difference was observed under static culture conditions. The absence of detectable differences in metabolic substrate utilization between human heart chambers may be attributed to plate-to-plate variability, which is difficult to avoid in human heart organotypic slice culture due to donor availability. Additionally, underlying comorbidities may have influenced the metabolic activity of the donor hearts (Table 2).

**Table 2.** Patient Demographics from Donor Human Hearts.

| Heart | Sex | Age (yrs) | BMI | Comorbidities | EF (%) | COD |
| --- | --- | --- | --- | --- | --- | --- |
| 1 | F | 59 | 29.5 | Sinus Tachycardia, LV Hypertrophy, Arthritis | NA | CVA/Stroke |
|  | F | 62 | 24.4 | Hypertension | NA | CVA/Stroke |
| 3 | F | 51 | 31.2 | Marijuana, Cigarettes, Cirrhosis | NA | CVA/Stroke |
| 4 | M | 58 | 23.3 | ALS | NA | Anoxia |
| 5 | F | 32 | 47.8 | Cigarettes, High Blood Pressure | 56 | CVA/Stroke |
| 6 | F | 48 | 33.3 | Hypertension | 75 | CVA/Stroke |

## NONSTANDARD ABBREVIATIONS AND ACRONYMS

ATP: Adenosine Triphosphate
BSA: Bovine Serum Albumin
BDM: 2,3-Butanedione monoxime
CPT1A: Carnitine Palmitoyl-Transferase 1A
DMSO: Dimethyl Sulfoxide
DPBS: Dulbecco’s Phosphate-Buffered Saline
ECAR: Extracellular Acidification Rate
EDTA: Ethylenediaminetetraacetic Acid
FA: Fatty Acid
FBS: Fetal Bovine Serum
GLC: Glucose
GLN: Glutamine
IC50: Half-Maximal Inhibitory Concentration
LA: Left Atrium
LV: Left Ventricle
NMR: Nuclear Magnetic Resonance
OCR: Oxygen Consumption Rate
PET: Photon Emission Tomography
RA: Right Atrium
RV: Right Ventricle
RVOT: Right Ventricular Outflow Tract
SERCA2a: Sarcoplasmic Reticulum Ca^2+^-ATPase
SPECT: Single-Photon Emission Computed Tomography
TCA: Tricarboxylic Acid

## Funding

This work is supported by American Heart Association (AHA) Sudden Cardiac Death SFRN Grant 19SFRN34830033 (to I.R.E.).

We gratefully acknowledge the families of organ donors and the staff at Washington Regional Transplant Community and Gift of Hope for their support in the procurement of human heart tissue. We also thank Paloma De Melo Amaral for technical support.

